# B cell repertoire sequencing of HIV-1 pediatric elite-neutralizers identifies multiple broadly neutralizing antibody clonotypes

**DOI:** 10.1101/2023.07.07.548149

**Authors:** Sanjeev Kumar, Prashant Bajpai, Collin Joyce, Sushil Kumar Kabra, Rakesh Lodha, Dennis R. Burton, Bryan Briney, Kalpana Luthra

**Author notes:** These authors contributed equally (Sanjeev Kumar, Prashant Bajpai).

## Abstract

A limited subset of HIV-1 infected adult individuals typically after at least 2-3 years of chronic infection, develop broadly neutralizing antibodies (bnAbs), suggesting that highly conserved neutralizing epitopes on the HIV-1 envelope glycoprotein are difficult for B cell receptors to effectively target, during natural infection. Recent studies have shown the evolution of bnAbs in HIV-1 infected infants. We used bulk BCR sequencing (BCR-seq) to profile the B cell receptors from longitudinal samples (3 time points) collected from a rare pair of antiretroviral-naïve, HIV-1 infected pediatric monozygotic twins (AIIMS_329 and AIIMS_330) who displayed elite plasma neutralizing activity against HIV-1. BCR-seq of both twins revealed convergent antibody characteristics including V-gene use, CDRH3 lengths and somatic hypermutation (SHM). Further, antibody clonotypes with genetic features similar to highly potent bnAbs isolated from adults showed ongoing development in donor AIIMS_330 but not in AIIMS_329, corroborating our earlier findings based on plasma bnAbs responses. An increase in SHM was observed in sequences of the IgA isotype from AIIMS_330. This study suggests that children living with chronic HIV-1 can develop clonotypes of HIV-1 bnAbs against multiple envelope epitopes similar to those isolated from adults, highlighting that such B cells could be steered to elicit bnAbs responses through vaccines aimed to induce bnAbs against HIV-1 in a broad range of people including children.

## INTRODUCTION

Human immunodeficiency virus type 1 (HIV-1) infections are a global health problem that has affected an estimated 38.4 million people worldwide including children(1). The integration of virus into the host cell genome and tremendous level of viral mutation observed in individuals living with HIV-1 are major obstacles to developing effective therapeutics and vaccines againstHIV-1(2,3). During HIV-1 infection, neutralizing antibodies (nAbs) and non-nAbs are elicited against multiple epitopes of the HIV-1 envelope glycoprotein(4–6). In the past 15 years, the invention of high-throughput technologies for antibody generation i.e. single B cell sorting, high-throughput antibody screening by micro-neutralization methods and single cell sequencing, have led to the discovery and molecular characterization of highly potent second-generation HIV-1 broadly neutralizing antibodies (bnAbs) both from adults and children living with HIV-1(5–7). Presently, researchers around the globe are leveraging our understanding of the genetic, functional, and structural properties of these bnAbs to develop HIV-1 vaccines that reliably induce broad and potently neutralizing antibody responses(8,9).

The characteristic features of HIV-1 bnAbs targeting multiple envelope epitopes that are isolated from adult donors have been extensively studied and are well-known(5,6). However, the understanding of genetic and molecular features of HIV-1 bnAbs from children is still limited. So far, only two HIV-1 pediatric bnAbs have been identified, BF520.1 by Simonich et.al.(10) and AIIMS-P01 by us (Kumar et. al.)(11). Further insights into the pediatric B cell repertoire are urgently needed to support next-generation vaccine design and vaccination strategies aiming to elicit protective bnAbs responses in a wide range of individuals including children. It has been reported in multiple studies that children living with HIV-1 showed broader and more potent bnAbs responses with multiple epitope specificities as compared to bnAbs from adults even within one-year of age, suggesting that bnAb responses in children are developed from different maturation mechanisms/pathways(10–21). This is supported by the discovery of BF520.1 and AIIMS-P01:pediatric bnAbs which exhibit comparable HIV-1 neutralization breadth and potency to adult bnAbs but with limited somatic hypermutation (SHM)(10,11).

In the past 15 years, we have established a rare cohort of HIV-1 clade C chronically infected pediatric donors including infants(11,14,16,17,21–25). Recently, we reported the identification of infant and adolescent pediatric elite-neutralizers (AIIMS_329 and AIIMS_330) from characterization of their plasma HIV-1 bnAbs responses at a single and longitudinal time points, respectively(14,21). We observed the development of bnAbs targeting the V1/V2 apex, glycan supersite, andCD4 binding site (CD4bs) in AIIMS_330. We also noted V1/V2 apex- and glycan supersite-dependent bnAbs in AIIMS_329, but at a lower potency and breadth in comparison to AIIMS_330(21). Herein, we performed deep sequencing of the bulk B cell repertoires (BCR-seq) of two monozygotic twin HIV-1 pediatric elite-neutralizers (AIIMS_329 and AIIMS_330). BCR-seq of both AIIMS_329 and AIIMS_330 showed convergent antibody characteristics including Variable gene use, heavy chain complementarity determining region (HCDR3) lengths, SHM frequency. Mapping BCR-seq data to knownHIV-1 bnAbs, allowed us to identify antibody clonotypes with similar features to potent HIV-1 bnAbs in AIIMS_330 but not in AIIMS_329, corroborating with our previous serological findings. This study is an important step toward defining specific features of the antibody repertoire and shared clonotype maturation that are associated with the development of HIV-1 elite-neutralizing activity. Some of the shared lineages identified in this pair of identical twins, who had acquired HIV-1 infection by vertical transmission, may have been elicited by exposure to common HIV-1 antigens. Defining the sequences of such shared clonotypes in a large number of children can shed light on the role of specific B cell receptor features in the response to HIV-1 infection.

## MATERIALS AND METHODS

### Patients Characteristics and Ethics Statement

Monozygotic twin antiretroviral naïve HIV-1 clade C pediatric elite-neutralizers (AIIMS_329 and AIIMS_330) were recruited from the Outdoor Patient Department of the Department of Pediatrics, All India Institute of Medical Sciences (AIIMS), New Delhi, India for this study at the age of 9 years and were followed for a total period of 60 months. Blood was drawn in 5-ml EDTA vials, and plasma was aliquoted and stored for plasma antibody-based HIV-1 neutralization assays, viral RNA isolation, and viral load determinations. The study was approved by the Institute Ethics Committee (IEC/59/08.01.16, IEC/NP-536/04.11.2013, IEC/NP-295/2011 and RP-15/2011). All experiments were conducted in accordance with the Institutional guidelines and protocols.

### Next-generation sequencing of HIV-1 pediatric B cell antibody repertoires

The deep sequencing of bulk BCR was performed using primers and protocols as described previously by Briney et.al.(38) Briefly, total RNA from 2 or 3 million peripheral blood mononuclear cells (PBMCs) was extracted (RNeasy Maxi Kit, Qiagen) from each time point (2015 (112 months p.i), 2016 (117 months p.i) and 2018 (138 months p.i)) and antibody sequences were amplified using methods and primers as previously described. The PCR product sizes were verified on agarose gel (E-Gel EX; Invitrogen) and quantified with fluorometry (Qubit; Life Technologies), pooled at approximately equimolar concentrations and each sample pool was re-quantified before sequencing on an Illumina MiSeq (MiSeq Reagent Kit v3, 600-cycle).

### Processing of next-generation sequencing data

The Abstar analysis pipeline was used as previously described to quality trim, remove adapters and merge paired sequences(38). Sequences were then annotated with Abstar in combination with UMI based error correction by AbCorrect (https://github.com/briney/abtools/). For comparison of frequencies, read counts were scaled for each repertoire as previously described due to the large differences in the number of reads between each group. Somatic hypermutation (SHM) was calculated using the R package Shazam(44).

### Clonotype analysis

Clonotype analysis was performed using Immcantation pipeline(44,45). Sequences were grouped into clonotypes based on nucleotide hamming distance of 0.16 calculated based on bimodal distribution of distance of each sequence with its nearest neighbor. Alternatively, sequences were also clustered into clonal groups using an in-house script. The criteria used for clonal assignment was sequences having same V and J gene usage, same CDRH3 length and at least 80% CDRH3 amino acid identity. Germline V(D)J sequence was reconstructed using IMGT-gapped reference V, D and J sequences.

### Deep mapping of HIV-1 mAbs to B cell repertoire sequencing data

List of HIV-1mAbs which have been reported previously was compiled from CATNAP database and literature(5–7,42). Antibodies for which the gene usage information and CDRH3 amino acid sequence was available were selected for downstream analysis. The list of selected mAbs is given in **Supplementary Table 7**. Each mAb was mapped to all the sequences to identify the ones which have the same V and J gene usage, same CDRH3 length and at least 50% identity in CDRH3 amino acid sequence.

### Alignment and phylogeny

Alignment was performed using R package MSA. Sequence logos were made using R package ggseqlogo. Distance between sequences were calculated using the neighbor joining method. Phylogenetic tree was constructed using R package ape.

### Statistical analysis

All analysis was performed using R programming language (version 4.1.0). Statistical significance between groups was estimated using Wilcox rank sum test. The following packages were used for the analyses: scatterpie (0.1.7), nortest (1.0-4), scales (1.1.1), ggrepel (0.9.1), stringr (1.4.0), tigger (1.0.0), ComplexUpset (1.3.0), ggpubr (0.4.0), alakazam (1.1.0), shazam (1.1.0), patchwork (1.1.1), writexl (1.4.0), readxl (1.3.1), dplyr (1.0.6), reshape2 (1.4.4) and ggplot2 (3.3.5).

## DATA AND CODE AVAILABILITY STATEMENT

Raw sequence data that support the findings in this study are available at the NCBI Sequencing Read Archive (www.ncbi.nlm.nih.gov/sra) under BioProjectnumber: PRJNA999025. Processed datasets are available at https://github.com/prashantbajpai/HIV_BCR_Analysis. All bash and R scripts required for reproducing the data can be found on github repository (https://github.com/prashantbajpai/HIV_BCR_Analysis). Raw data used for generating the figures are available at the github page or in the supplementary files.

## RESULTS

### Deep BCR sequencing of HIV-1 pediatric elite-neutralizers

AIIMS_329 and AIIMS_330 are monozygotic twins, and both display elite HIV-1 plasma neutralization. Longitudinal samples were obtained from both twins prior to their initiation of combined antiretroviral therapy (cART) in 2018. The total number of sequences obtained for each subject after sequencing and annotation ranged from approximately 1.0×10^5^ to 4.0×10^5^ sequences (**Supplementary Table 1**). Immunogenetic analysis of heavy chain Variable (VH) genes identified IGHV3-21, IGHV3-23, and IGHV-34 to be predominant gene usages in both AIIMS_329 and AIIMS_330 groups in the three timepoints of their sample collection in the years 2015 (112 months p.i.), 2016 (117 months p.i.), and 2018 (138 months p.i.). Within IGHV3-21, AIIMS_330 from the three timepoints showed a trend towards higher frequency compared to AIIMS_329, and 330_2015 had the highest frequency (25.6% compared to compared to approx. 13 other groups, **Figure 1A, Supplementary Table 2**). No major differences in the VH gene frequency of the two groups-at any time point was observed (**Figure 1A, Supplementary Table 2**). Consistent with previous studies, IGHD3-22, IGHD3-10, IGHD6-13 were found to be major D genes and IGHJ4 was found to be predominant heavy J gene. No noteworthy differences were observed in the frequencies of heavy chain Diversity (D)or Joining (J) genes (**Figure 1B and IC**). In the light chain, IGKV4-1 and IGKV3-20 were the predominant gene usages. We observed a trend towards higher frequency of IGKV4-1 in AIIMS_330 compared to AIIMS_329 (14.54%, 20.35%, 14.16% in 330_2015, 330_2016 and 330_2018 compared to 8.89%, 10.27%, 12.99% in 329_2015, 329_2016 and 329_2018). We also observed frequency of IGLV2-14 to be lower in AIIMS_330 compared to AIIMS_329 (2%, 2.09%, 2.13% in 330_2015, 330_2016 and 330_2018 compared to 4.63%, 7.91%, 5.09% in 329_2015, 329_2016 and 329_2018) (**Figure 1D, Supplementary Table 3**). We also observed marked differences in the light chain J gene use between the two groups. Gene usage IGKJ2 showed a trend towards higher frequency in AIIMS_330 at three timepoints. Further, the frequency of IGLJ3 was markedly lower in AIIMS_330 compared to AIIMS_329.Overall, though heavy chain V, D, and J gene frequencies were similar in AIIMS_329 and AIIMS_330, we observed marked differences in lambda and kappa light chains between the two groups at all timepoints.

**Figure 1:**
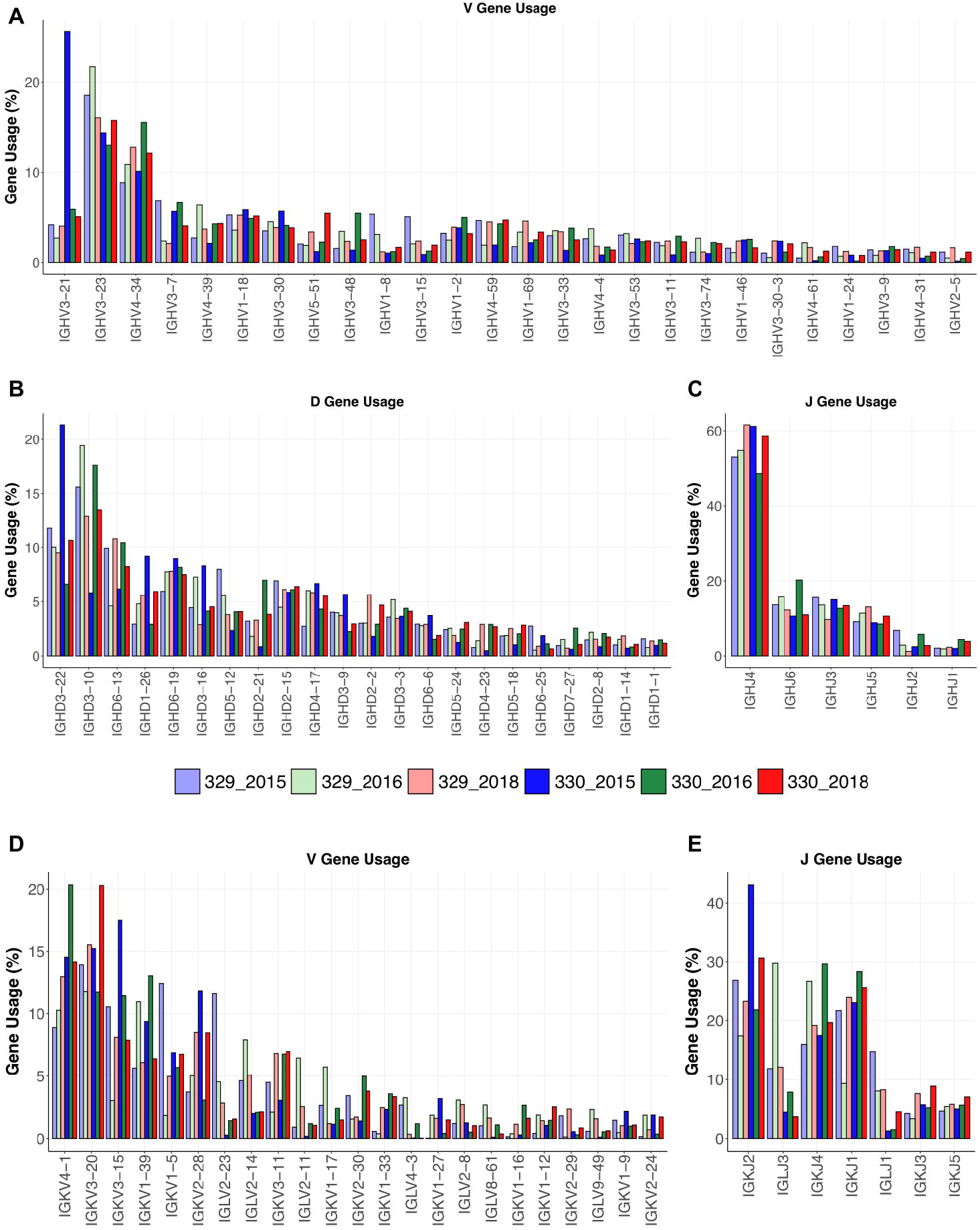
Gene usage of heavy and light chains. V-gene **(A)**, D-gene **(B)** and J-gene **(C)** usage of heavy chains of genes shown as percentage of total sequences and arranged by descending frequency. V-gene **(E)** and D-gene **(F)** of light chains are shown.

### Shared immunogenetic features between AIIMS_329 and AIIMS_330

The CDRH3 length distributions were similar for AIIMS_329 and AIIMS_330 at all time points, with mean CDHR3 length being lower in the AIIMS_330(12.91, 13.22 and 12.66 in 330_2015, 330_2016, and 330_2018, respectively, compared to 12.97, 13.37, and 12.97 in 329_2015, 329_2016, and 329_2018) (**Figure 2A, Supplementary Table 4**). A similar trend was observed in the distribution of light chain CDR3 lengths (**Supplementary Fig 1**). SHM frequency was significantly higher in AIIMS_330 at all three timepoints compared to AIIMS_329 group. In the 2018 timepoint, median SHM in AIIMS_330 was 2.42% compared to 0.41% in AIIMS_329 (**Figure 2B, Supplementary Table 5**). Interestingly, when the sequences were segregated into isotypes, sequences from IgA isotype were observed to have significantly higher SHM in AIIMS_330 compared to AIIMS_329. The median SHM in IgA sequences was found to be 11.74%, 6.97%, and 6.94% in 330_2015, 330_2016, and 330_2018, respectively, compared to 7.63%, 6.05%, and 4.72% in 329_2015, 329_2016, and 329_2018. No significant changes were observed in other isotypes (**Figure 2C, Supplementary Table 6**).

**Figure 2:**
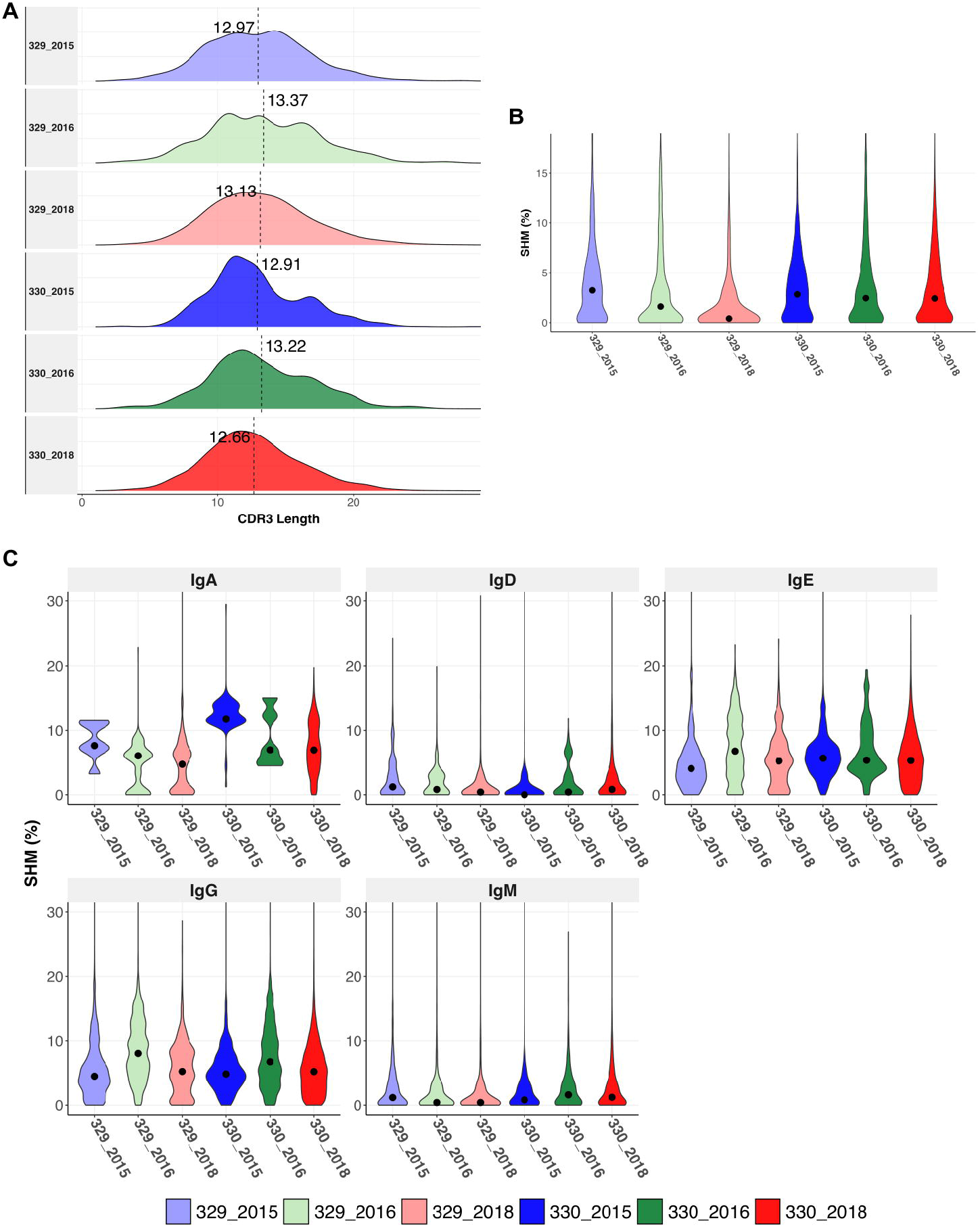
Immunogenetics characteristics of heavy chain sequences. **(A)** CDRH3 length distribution of heavy chains in each group. The dotted horizontal line shows the median of each group. **(B)** Violin plot shows somatic hypermutation (SHM) in each group. The dot represents the median in each group. **(C)** Violin plot shows the SHM in each group segregated by isotype. The dot represents the mean in each group.

### Shared clonotype analysis identified distinct features between AIIMS_330 and AIIMS_329

Sequences were subsampled before performing the clonotype analysis (more detail of the subsetting approach can be found in the methods section). Frequency of total shared clonotypes between any two groups was found to be less than 0.07% (77 shared clonotypes across 10918 total clonotypes, **Figure 3A bottom panel**). Of all the clonotypes identified in each group, clones in 330_2018 timepoint had the highest SHM compared to others. Most of the clonotypes in 330_2018 timepoint had mean SHM >10%. We also observed higher expansion in 2018 timepoint in AIIMS_330 compared to AIIMS_329 as observed from lower number of clonotypes and higher number of sequences in each clonotype. Interestingly, the overall clonotypes in 2018 timepoint in both groups were higher compared to 2015 and 2016 timepoints (**Figure 3A middle panels**). No notable differences were observed in the V-family distribution of shared clonotypes (**Figure 3A top panel**). We also analyzed the clones shared between each sample and timepoints. We did not observe any clonotypes shared between all the groups or even between AIIMS_330 and AIIMS_329 at all three timepoints. Interestingly, the highest number of shared clonotypes were identified between 330_2016 and 330_2018 timepoints (39 shared clonotypes) but not between any timepoints in AIIMS_329. Of these 39 clonotypes, only 3 had mean SHM greater than 10%. However, those 3 clonotypes did not have any significant differences in their CDR3 amino acid (**Figure 3B**). We identified 12 clonotypes that were shared between AIIMS_330 and AIIMS_329 at 2016 timepoint of which 5 clonotypes had mean SHM greater than 10%. These 5 clonotypes were found to have accumulated higher mutations in CDR3 compared to the 3 clonotypes shared between 330_2016 and 330_2018 (**Figure 3C**). None of these shared clonotypes were predicted to encode sequence insertions or deletions which are frequently found in HIV bnAbs and thought to be difficult to elicit by vaccination. Such BCRs could plausibly define a more efficient path for bnAb maturation that can be exploited by rational HIV vaccine immunogens. We also observed 21 clonotypes in common between 2018 timepoint between AIIMS_330 and AIIMS_329. Of these, only one clonotype had >10% SHM. CDR3 of sequences from these clonotypes were observed to harbor higher number of mutations in 330_2018 timepoint compared to 329_2018 (**Figure 3D**).

**Figure 3:**
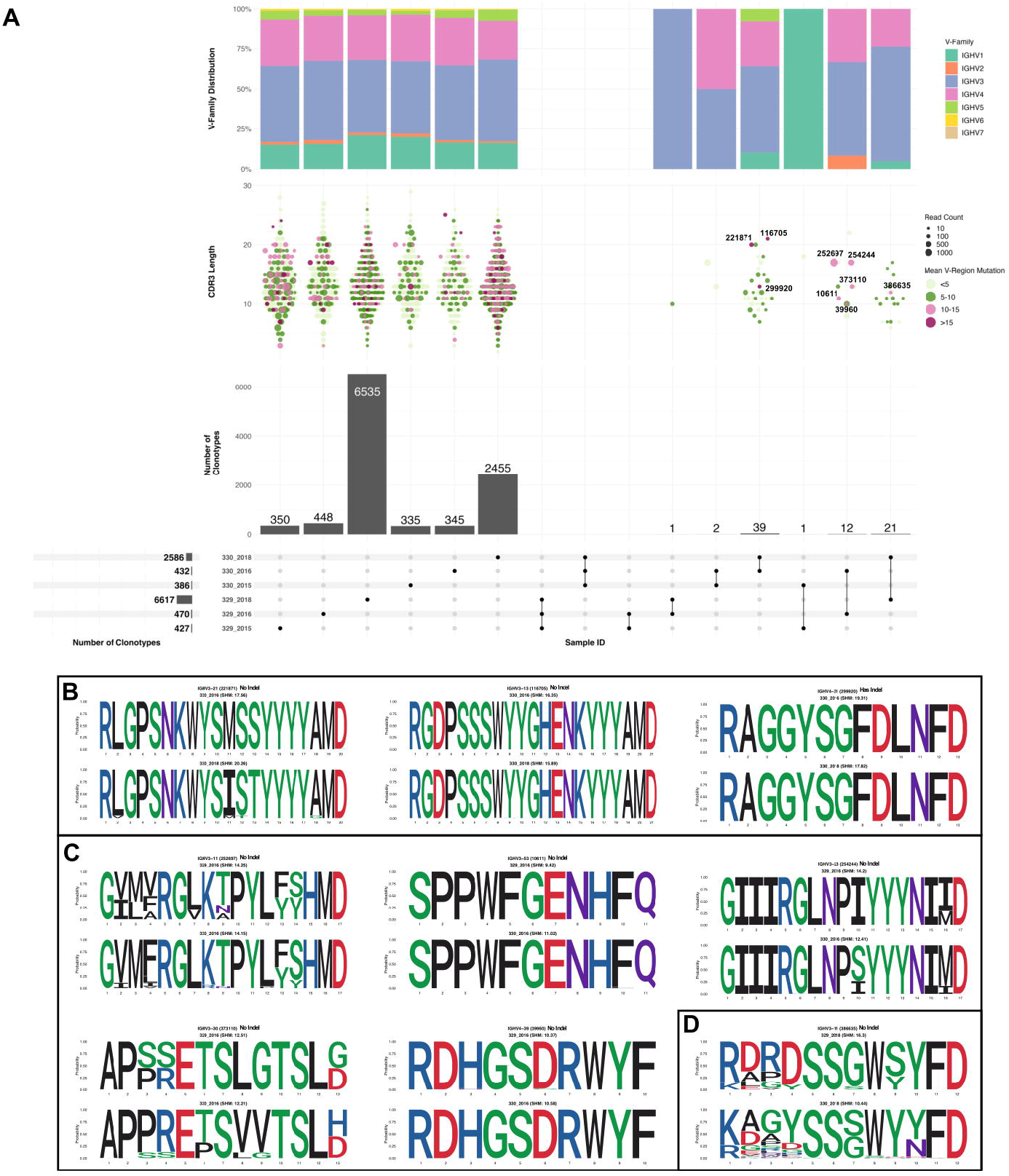
Shared clonotypes between the groups. **(A)** Upset plot illustrating the intersection of clonotypes between the subject 329 and 330 at the three time points (2015, 2016 and 2018). Only the clonotypes which were not singlets with more than one sequences were selected for the analysis. Each vertical bar shows the number of clonotypes present in each group indicated by the intersection matrix below it (where a single dot in the matrix is a single group). The number above the bars show the number of common clonotypes between the groups of intersecting sets of clonotypes marked below the bars. For each intersection the dot plot above indicates the CDRH3 length distribution for each clonotype. Size of each dot indicates the number of sequences present in each clonotype and the color shows the somatic hypermutation (SHM) frequency. The vertical bar above it shows the distribution of VH genes for each intersection. Within the shared clonotypes, the clones with SHM >= 10% are labelled and shown. **(B)** Sequence logo representing alignment of CDRH3 region of clones common between 2016 and 2018 timepoint of subject 330 with >= 10% SHM. **(C)** Sequence logo representing alignment of CDRH3 region of clones common between 2016 and subject 329 and 330 with >= 10% SHM. **(D)** Sequence logo representing alignment of CDRH3 region of clones common between 2018 timepoint of subject 329 and 330 with >= 10% SHM.

### Mapping of sequences to published adult HIV bnAbs identified 330_2018 to be capable of producing bnAbs with characteristics of adult bnAbs

To identify if either of the two children could generate known HIV-1 bnAbs at any timepoint, we mapped the sequences from each group to known HIV mAbs compiled from HIV neutralizing antibody database (CATNAP) and published studies. A total of 347 mAbs were compiled (**Supplementary Table 7**) and mapped to sequences from each group. The criteria for calling a sequence a successful hit is described in the methods. Of the total 347 mAbs, 78 mAbs mapped to at least one of the sequences in at least one of the groups. Surprisingly, majority of the sequences that mapped to these mAbs were from 330_2018 timepoint but no other timepoints. Moreover, we observed the expansion/more frequency of the following HIV-1 bnAbs like sequences in AIIMS_330:VRC38.01 (V2 apex)(26), VRC34.01 (fusion peptide)(27), HGN194 (V3)(28), HK20 (gp41 HR)(29), DH270 (glycan supersite)(30), BG505.m27 (V3)(31), 8ANC131 (CD4bs)(32), 2558 (V3)(33), and VRC33.01 (CD4bs)(34) (**Figure 4A**). Strikingly, except DH511.1, DH511.6 (MPER)(35)and BG505.m27 HIV-1 bnAbs, no other mapped sequences from AIIMS_330 only showed SHM >10%, however, such high SHMs were not observed for any of the mapped sequences from AIIMS_329. The 78 mAbs that were mapped to heavy chain sequences were also mapped to the light chain sequences. Using the same mapping criteria, it was found that 28 of the mAbs also mapped to light chain sequences and sequences from AIIMS_330 showed higher SHM compared to other timepoints (**Figure 4B-C**). In summary, we show that both heavy and light chain sequences mapped to similar set of mAbs and showed higher SHM and number in AIIMS_330.

**Figure 4:**
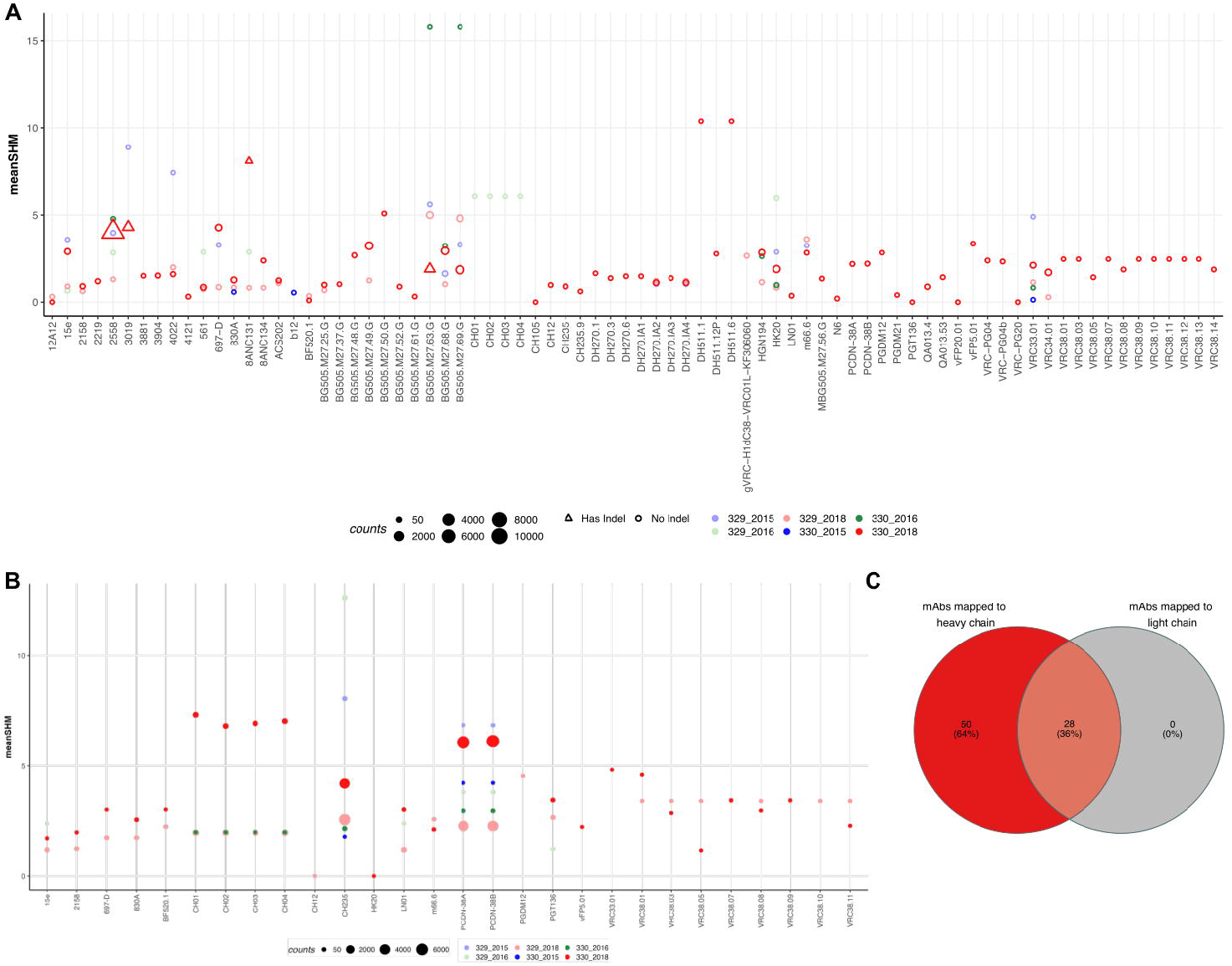
Sequences shared with publicly available HIV bnAbs. Publicly available mAbs mapped to the heavy chain **(A)** and light chain **(B)** sequences are shown. Each HIV mAb is shown on the x-axis while and each dot represents the number of sequences mapped to the respective mAb. The criteria for calling mAb a hit was same V and J gene, more that 50% identity in the CDHR3 amino acid and same CDRH3 length. The size of the dot represents the number of sequences mapped and color represents the group. On y-axis mean somatic hypermutation (SHM) is shown. **(C)** Venn diagram showing the overlap of mAbs that mapped in both heavy and light chain sequences.

## DISCUSSION

An understanding of pediatric HIV-1 bnAb responses elicited during chronic infection can provide critical insights for the design and development of effective universal HIV-1 vaccines for both adults and children capable of eliciting potent HIV-1 bnAb responses(36,37). Human antibody repertoire comprises a whole group of antibody genes generated in an individual since birth, which can be specific to an antigen during an infection/disease and vaccination. BCR-seq is a powerful method that has enabled the in-depth analysis of genetic features of antibody responses and tracking of antibody evolution during an infection/vaccination(38). Studying such antibody repertoires could reveal important information that include germline antibody genes, junctional diversity, SHM, clonotypes and allows identification of rare antibody lineage genes specific to a particular antigen.

Recently, in a study based on longitudinal characterization of plasma samples we reported the identification of adolescent pediatric elite-neutralizers who were HAART-naïve and living with chronic HIV-1 clade-C infection for more than 11 years(21). We evaluated the V1/V2 apex, glycan supersite, and CD4bs specific responses in AIIMS_329 and AIIMS_330, using ELISA and point mutated HIV-1 pseudoviruses based neutralization assays(21). We showed a longitudinal development of HIV-1 bnAbs targeting V1V2, N332 and CD4bs in AIIMS_330, whereas V1V2 and N332-supersite dependent HIV-1 bnAbs in AIIMS_329, but at a lower potency and breadth in comparison to AIIMS_330(21). Herein, we performed bulk BCR-seq of these two monozygotic twins HIV-1 pediatric elite-neutralizers (AIIMS_329 and AIIMS_330) to understand their BCR repertoire. Analysis of three time points (112, 117 and 138 months p.i.) from these pediatric elite-neutralizers showed high convergence of antibody gene-usage, CDRH3 lengths, SHM, and clonotypes. Though a higher IgA SHM was observed in AIIMS_330, no other isotype showed differential mutation patterns. The development of HIV-specific IgA responses with affinity maturation have been shown to in association with anti-gp41 IgA antibodies that occurs to a greater extent in elite controllers than in individuals on HAART(39), suggesting IgA in AIIMS_330 could be associated with lower viral load as compared to AIIMS_329.

Current common strategies to generate antigen specific antibodies and evaluating humoral vaccine responses are: 1) high-throughput sorting antigen-specific B cells followed by paired heavy and light chain gene amplification and sequencing, 2) barcoded antigen approaches like LIBRA-seq which leverage single cell microfluidics to link antibody genetics and binding specificity in high throughput, and 3) activation and culture of primary B cells followed by functional supernatant screening to identify B cell clones with desirable functional profiles. Such methods are labor-and time-intensive ways of identifying antigen-specific antibodies. Mapping sequences to well-known established antibody CDR3 information can provide templates or blueprints to identify important antibody genes in a distinct set of individual populations, antibody discovery, and vaccine response evaluation without the need for antigen-specific sorting(40). Similar methodologies have been used successfully to identify potent antibodies against Dengue, HIV-1, SARS-CoV-2, and influenza(41).

Here, we mapped BCR sequencing data against datasets of known HIV-1 bnAbs(42). Interestingly, this analysis led to the identification of multiple HIV-1 specific antibody clonotypes isolated from both adults and children in our BCR-seq data. AIIMS_330 showed clonotypes similar to several HIV-1 bnAbs identified in adults which are known to target multiple epitopes including MPER, CD4bs, N332, FP and V1V2, suggesting that antibody data mining based on CDRH3 sequences could help in bnAbs lineage identification to a particular antigen, as recently observed by an AI-based pipeline developed by Wu et.al.(41) for COVID-19 and Flu antibody genes and DENV specific mAbs by Durham et.al.(43) Our analysis of these bnAb lineages showed the evolution/more frequency (**Figure 4**) of these antibodies in AIIMS_330 donor, which corroborates with our previous findings based on plasma characterization and evolution of multi-epitope specific antibodies including CD4bs bnAbs in AIIMS_330 than AIIMS_329(21). Overall, the observation of some the shared lineages in this pair of identical pediatric twins, who had acquired HIV-1 infection by vertical transmission, suggest that they could evolve in response to common antigenic stimulus. Defining the sequences of such shared clonotypes, in a large number of children living with chronic HIV-1 infection worldwide, can shed light in understanding the role of specific B cell receptor repertoires in HIV-1 infection.

Our study provides insight into the BCR repertoires of pediatric twins capable of exceptionally broad and potent HIV-1 neutralization. We noted the presence of clonotypes with genetic similarity to known adults’ HIV-1 bnAbs targeting multiple epitope specificities in children, which corroborated the results of our previous study which characterized the plasma neutralization breadth, potency, and epitope specificity from the same pair of donors(21). We anticipate this data will be useful for the design and development of effective vaccine candidates and strategies for both adults and children to combat HIV-1. Our BCR-seq findings from this unique pair of pediatric elite-neutralizers suggests that multivalent HIV-1 vaccine development strategies focused on inducing a diverse range of HIV-1 bnAb specificities is likely superior to approaches that focus solely on a single epitope.

## ACKNOWLEDGEMENTS

We are thankful to the SERB, India (CRG/2021/003984), and Department of Biotechnology (DBT) (BT/PR 24520/MED/29/1222/2017) for the funding provided to K.L. This study is supported in part by ViiV Healthcare and the HIV Research Trust 2019 awarded to S.K. as part of his Graduate (Ph.D.) work training and gaining expertise in Next Generation Sequencing based technologies to sequence single B cell and bulk B cell repertoire sequencing. The funders had no role in the decision to publish or the preparation of the manuscript.

## AUTHOR CONTRIBUTIONS

Conceptualization: K.L., B.B., and S.K.; Methodology: B.B., C.C.J., and S.K.; Software: B.B., C.C.J., P.B.; Validation: B.B.; Formal analysis: S.K., and P.B.; Investigation: S.K., and C.C.J.; Resources: D.R.B., R.L, and S.K.K.; Data Curation: B.B., P.B., and C.C.J.; Management: S.K., P.B., and B.B.; Writing - Original Draft: S.K., and P.B.,; Writing - Review & Editing: K.L., B.B., and D.R.B.; Visualization: S.K., and P.B.; Supervision: K.L., B.B. and D.R.B.; Project administration: S.K.; Funding acquisition: K.L., B.B., and S.K.

## DECLARATION OF INTERESTS

All other authors declare no competing interests.

## Notes

### Competing Interest Statement

The authors have declared no competing interest.

### Summary of Updates

This version is updated with text, Figure 4.

