## Supplementary material for "B cell repertoire sequencing of HIV-1 pediatric elite-neutralizers identifies multiple broadly neutralizing antibody clonotypes": Supplementary File.pdf

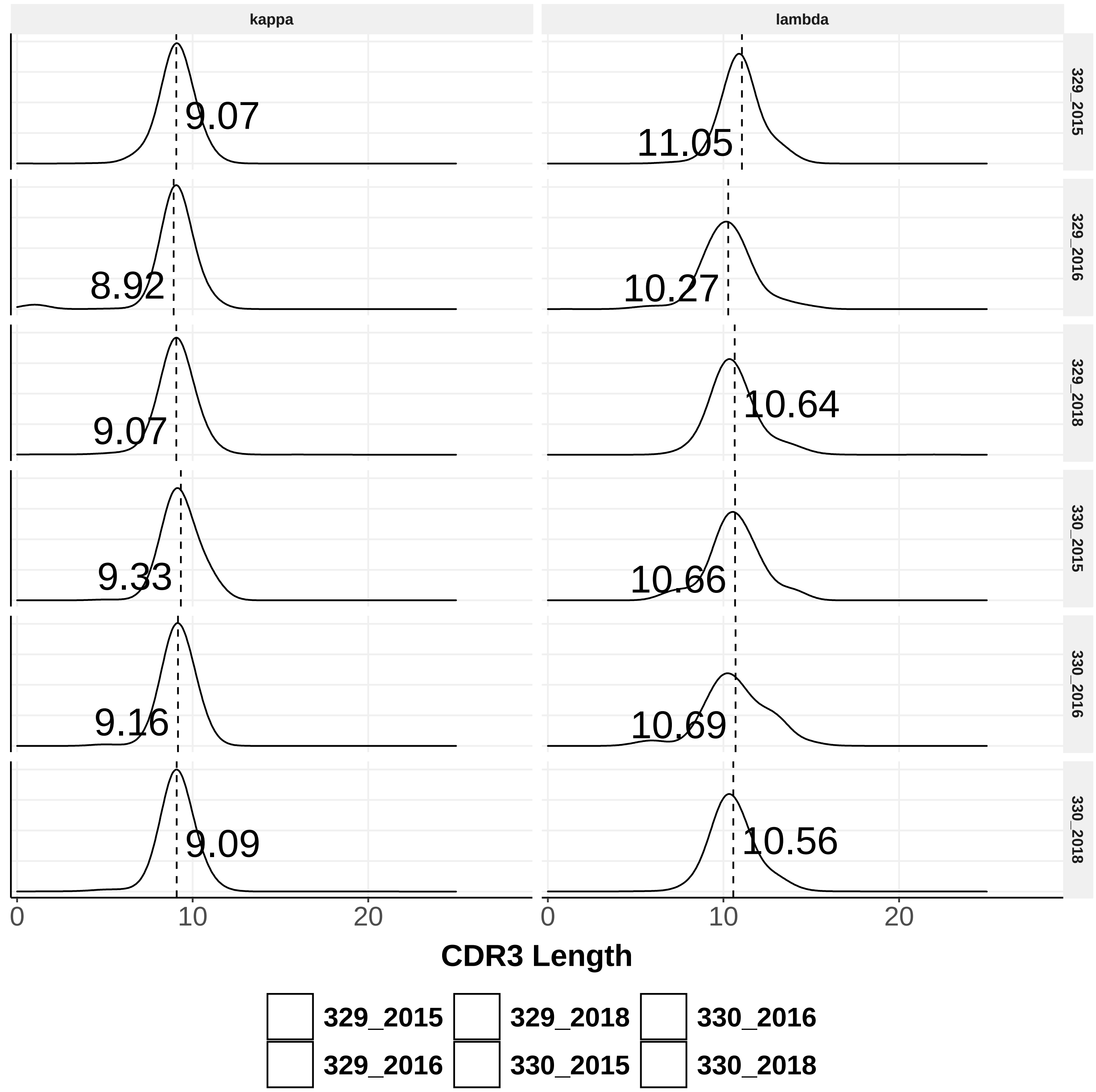


**Supplementary Figure 1**. CDRH3 length distribution of kappa and light chains in each group. The dotted horizontal line shows the median of each group.

**Supplementary table 1**: Number of sequences obtained from each sample post annotation with abstar pipeline.

| Sample ID | Number of sequences post- annotation |
|---|---|
| 329_2015 | 97888 |
| 329_2016 | 86178 |
| 329_2018 | 219087 |
| 330_2015 | 106076 |
| 330_2016 | 29255 |
| 330_2018 | 400249 |
